# An ethnobotanical study of the genus *Elymus*

**DOI:** 10.1101/734525

**Authors:** Emma S. Frawley, Claudia Ciotir, Brooke Micke, Matthew J. Rubin, Allison J. Miller

## Abstract

Grains of domesticated grasses (Poaceae) have long been a global food source and constitute the bulk of calories in the human diet. Recent efforts to establish more sustainable agricultural systems have focused in part on the development of herbaceous, perennial crops. Perennial plants have extensive root systems that stabilize soil and absorb water and nutrients at greater rates than their annual counterparts; consequently, perennial grasses are important potential candidates for grain domestication. While most contemporary grass domesticates consumed by humans are annual plants, there are over 7,000 perennial grass species that remain largely unexplored for domestication purposes. Documenting ethnobotanical uses of wild perennial grasses could aid in the evaluation of candidate species for *de novo* crop development. The objectives of this study are 1) to provide an ethnobotanical survey of the grass genus *Elymus*; and 2) to investigate floret size variation in species used by people. *Elymus* includes approximately 150 perennial species distributed in temperate and subtropical regions, of which at least 21 taxa have recorded nutritional, medicinal, and/or material uses. *Elymus* species used for food by humans warrant pre-breeding and future analyses to assess potential utility in perennial agricultural systems.

## Introduction

It is estimated that between 20% to 50% of the nearly 400,000 extant plant species in the world may be edible to humans (Füleky 2009; Warren 2015); however, only 6,000 of these have been cultivated for human consumption (FAO 2019). Cereals, members of the grass family (Poaceae), include several widely cultivated species, such as barley (*Hordeum vulgare* L.), maize (*Zea mays* L.), oats (*Avena sativa* L.), rice (*Oryza sativa* L.), rye (*Secale cereale* L.), sorghum (*Sorghum bicolor* L. Moench), sugarcane (*Saccharum officinarum* L.), and wheat (*Triticum aestivum* L.), among others (NGS 2008). Cereals are a staple of the human diet and comprise 50 percent of global caloric intake (Awika 2011; Warren 2015). Maize, rice, wheat, and sugarcane account for over half of the total crop production worldwide (FAO 2019), indicating their importance in the global food system and the relatively small number of grass species used in modern agriculture (e.g., Khoury et al. 2014).

Cereal domestication began at least 12,000 years ago and resulted in morphological and genetic changes in cultivated plants relative to their wild progenitors (Glémin and Bataillon 2009; Olsen 2013a; Olsen 2013b). For example, domesticated grass species exhibit a reduction in axillary branching, synchronization of maturation, and easy threshing (Zohary et al. 2012). Further, domesticated grasses have larger seeds that require reduced stratification and display decreased dormancy, shattering, and reduced or absent awns (Glémin and Bataillon 2009, Harlan et al. 1973; Harlan 1992). These characteristics contribute to more uniform harvest time, plants that can be grown in denser stands, increased seedling vigor, and more efficient harvesting (Glémin and Bataillon 2009). Subsequent crop improvement programs have focused largely on enhanced grain production and nutritional qualities of domesticated grasses, resulting in important alterations to a variety of seed traits, among other characteristics.

Grass species involved in early domestication processes were almost exclusively annuals (NGS 2008), perhaps due to their high seed output (Cox 2009), adaptation to early agricultural lands (DeHaan and Van Tassel 2014), and/or response to early selection efforts targeting synchronized maturation (Glémin and Bataillon 2009). However, ecological impacts of agricultural systems based on annual plants, including ongoing soil erosion and soil degradation (e.g. Montgomery 2007) have turned attention to the potential role of herbaceous, perennial species in contemporary agricultural systems. Perennials have deep root systems and longer growing seasons resulting in reduced erosion risk and greater plant productivity over time (Glover et al. 2010). Additionally, perennial species may be better adapted to temperature increases driven by climate change, as they are less affected by changes in the uppermost soil layer (Cox et al. 2006). As such, perennial crops may have an important role to play in the development of more sustainable agricultural systems (Bommarco et al. 2013; Cassman 1999; Ciotir et al. 2016; Ciotir et al. 2019; Cox et al. 2002; Doré et al. 2011; FAO 2009; Glover et al. 2010; Tittonell 2014).

Despite their potential utility, very few perennial grasses have been domesticated (Van Tassel et al. 2010). Several hypotheses have been proposed to explain the near absence of perennial, herbaceous crops. For example, some have suggested that their conservative resource allocation to reproductive structures relative to vegetative structures hinders response to selection for increased seed; others have proposed that herbaceous perennial plants exhibit reduced competitive ability in agricultural habitats compared to annual species (DeHaan et al. 2010; DeHaan and Van Tassel 2014). However, expanding understanding of agro-ecology, combined with new tools and analytical approaches, is driving increasing interest in pre-breeding of wild, herbaceous, perennial species. Several herbaceous, perennial species are currently under development, including perennial rice, sorghum, and wheat, among others (e.g., Cox et al. 2018; DeHaan et al. 2016; Huang et al. 2018).

There are two primary ways in which perennial grass crops can be developed (DeHaan and Van Tassel, 2014). First, annual crops can be hybridized with their perennial wild relatives. This serves to introgress annual traits (like high yield, abiotic stress tolerance) into a perennial background (e.g. perennial wheat (*Triticum aestivum* x *Thinopyrum intermedium*) (DeHaan et al. 2018; Hayes et al. 2018) or vice versa. A second means of developing perennial grass crops is through *de novo* domestication of wild species, as is underway, for example, with the wild wheat relative Kernza (*T. intermedium* (Host) Barkworth & D.R. Dewey) at the Land Institute (Salina, KS). However, one of the current challenges for *de novo* domestication is the identification of wild species for inclusion in pre-breeding programs (Ciotir et al. 2019).

When investigating wild plant species with potential utility in perennial agricultural systems it is valuable to consider historical and contemporary ethnobotanical uses, as well as their fundamental morphological features and geographic distributions. Ethnobotanical and other data on plant diversity and use, including records of plant form preserved in herbarium specimens, are often housed in botanical gardens and museums (Miller et al. 2015). These records offer a unique opportunity to explore agriculturally relevant questions about potential candidates for domestication. For example, within a particular genus of grasses, how many species are perennial? How many species have been used by people, what parts of the plant have been used, and for what purposes?

*Elymus* L. (wild rye) is an appealing genus for perennial grain domestication because of its compact and determinate inflorescence structure, capacity to self-pollinate, and current use as forage, among other characteristics. Several *Elymus* species have been developed as forage cultivars (e.g. blue wildrye (*E. glaucus* Buckley), thickspike wheatgrass (*E. lanceolatus* Scrib. & J.G. S.M), Canada wild rye (*E. canadensis* L.), slender wheatgrass (*E. trachycaulus* Link), Snake River wheatgrass (*E. wawawaiensis* J. Carlson & Buckley) and Virginia wildrye (*E. virginicus* L.) (Aubry et al. 2005; Lloyd-Reilley 2010; Tilley et al. 2011). To date, multiple *Elymus* species have been hybridized in a variety of pre-breeding initiatives. For example, there are at least seventeen *Elymus-*wheat hybrids (Cox et al. 2002) that have been developed for drought and salt tolerance (i.e. *Elymus mollis* Trin. x *Triticum durum* Desf.; Fatih 1983) and scab resistance (i.e. *E. trachycaulus* x *T. aestivum* L.*, E. tsukushiensis* Honda x *T. aestivum;* Kole 2011; Wang et al. 1999). Other *Elymus* hybrids include *Elymus hoffmannii* R.B. Jensen & R.H. Assay, an advanced generation hybrid between quackgrass (*E. repens* L.) and bluebunch wheatgrass (*Pseudoroegneria spicata* (Pursh) Á. Löve) with drought and salinity tolerance (St. John 2010). This work indicates *Elymus* is amenable to breeding processes and that some species within the genus may hold promise for perennial grain crop development.

In this study we investigate *Elymus* to provide information that might facilitate evaluation of species for use in *de novo* domestication processes. The specific objectives of this study were to: 1) conduct an ethnobotanical survey of the genus *Elymus*; and 2) investigate floret size in species used by people. These data provide valuable information about *Elymus* use and floret size variation, and underscore how ethnobotanical studies can aid agricultural processes through the evaluation of wild species and their potential applications in pre-breeding processes.

## Methods

### Study System

*Elymus* includes approximately 150 wild, herbaceous, perennial species distributed across North Temperate regions (Barkworth 2007; Lu 1993), including 39 species that occur in North America (32 of which are native; Barkworth 2007). *Elymus* caryopses (grains) are typically oblong to oblong-linear and adherent to the lemma and palea with hairy apices (Barkworth 2007; Chen and Zhu 2006; Lu 1993). Inflorescences are erect spikes with one to three spikelets at each node. Spikelets are ordinarily sessile with one to 11 florets. The lower florets are typically functional, and the distal florets are often reduced (Barkworth 2007; Chen and Zhu 2006; Kellogg 2015). Species that occur in western and northern North America have solitary spikelets, whereas those found east of the Rocky Mountains have multiple spikelets per node (Barkworth 2007).

### Inclusion of Leymus

Since the initial description of *Elymus* by Linnaeus, its taxonomy has varied under different taxonomic treatments (Helfgott and Mason-Gamer 2004; Lu 1993). Of particular interest to this study is the genus *Leymus* Hochst., whose species have often been included in circumscriptions of *Elymus*. Three *Leymus* species presented in this survey, *L. cinereus* (Scribn. & Merr.) Á. Löve*, L. condensatus* (J. Presl) Á. Löve, and *L. triticoides* (Buckley) Pilg., were originally described as *Elymus* species by previous authors (*E. cinereus* Scribin. & Merr., *L. condensatus* J. Presl, and *E. triticoides* Buckley), but are now considered synonyms for *Leymus.* We included these species in our results because some ethnobotanical descriptions surveyed here treat them as *Elymus*, and all three were used extensively by indigenous communities in the American southwest.

### Ethnobotanical analysis of Elymus

We performed a literature review to investigate recorded uses of *Elymus* species. We surveyed 121 print resources accessed at the Peter H. Raven Library at the Missouri Botanical Garden library. We reviewed 1) general ethnobotanical studies carried out in regions in which *Elymus* is known to occur; 2) ethnobotanical studies focused specifically on cultures of native communities located in these regions; and 3) global assessments of edible plants. We surveyed two online ethnobotany databases, *Native American Ethnobotany Database* (http://naeb.brit.org/) and *Plants for a Future* (https://pfaf.org/), and two online scientific databases, *JSTOR* (http://www.jstor.org) and *Web of Science* (http://www.webofknowledge.com/WOS) for relevant information about *Elymus.* We collected data on historical use by indigenous communities, human and animal edibility, cultivation history, and the uses of different plant parts. Results were recorded in the Perennial Agriculture Project Global Inventory online database (http://www.tropicos.org/Project/PAPGI). We also collected data on geographic distributions from specimen data at the Missouri Botanical Garden herbarium and from the Global Biodiversity Information Facility (www.gbif.org). Ethnobotanical uses were categorized as food, forage, medicine, and/or material. The food category included species that were consumed by humans; the forage category identified species cultivated for growth in pastures and for consumption by livestock; the medicine category designated species that were used in ceremonial decoctions or had therapeutic or healing utilities; finally, the material group covered species used as tools, housewares, and in construction, as well as other applications as raw materials.

### Measurements of floret traits from herbarium specimens

Grain morphology is an important target of selection in grass species undergoing domestication for human consumption (Glemín and Batallion 2009). While many wild species have relatively small, long, thin grains, selection during domestication generally favors larger, rounder grains (Gegas 2010; Okamoto 2012; Stougaard and Xue, 2004). We were interested in surveying grain size variation in species with documented ethnobotanical uses. We hypothesized that *Elymus* species used for human consumption may display larger grain sizes than those used for other purposes. A definition of the “pure seed unit” for crop conditioning is the floret: the reproductive structure including the lemma, palea, and caryopsis (grain), and excluding the awn when the awn length is longer than that of the entire floret (Gregg and Billups 2010). There is a positive correlation between floret cavity size (volume) and grain growth, including grain size and weight (Millet and Pinthus 1984; Millet 1986).

We calculated floret area for *Elymus* species with documented histories of use to examine relationships between floret size, ethnobotanical use, and collection location. Our ethnobotanical analysis identified 21 species with ethnobotanical uses (see results below). For each of these 21 species, we selected *Elymus* specimens from the herbarium at the Missouri Botanical Garden based on their collection location, targeting specimens that had been collected in a country or state where *Elymus* use by indigenous communities had been documented (Figure 1). If there was no indigenous community specifically identified for a taxon, we selected a specimen from the species known native range. For example, because *E. canadensis* was used historically in Utah and Colorado, sampled specimens came from these states (Table 1).

**Table 1.**
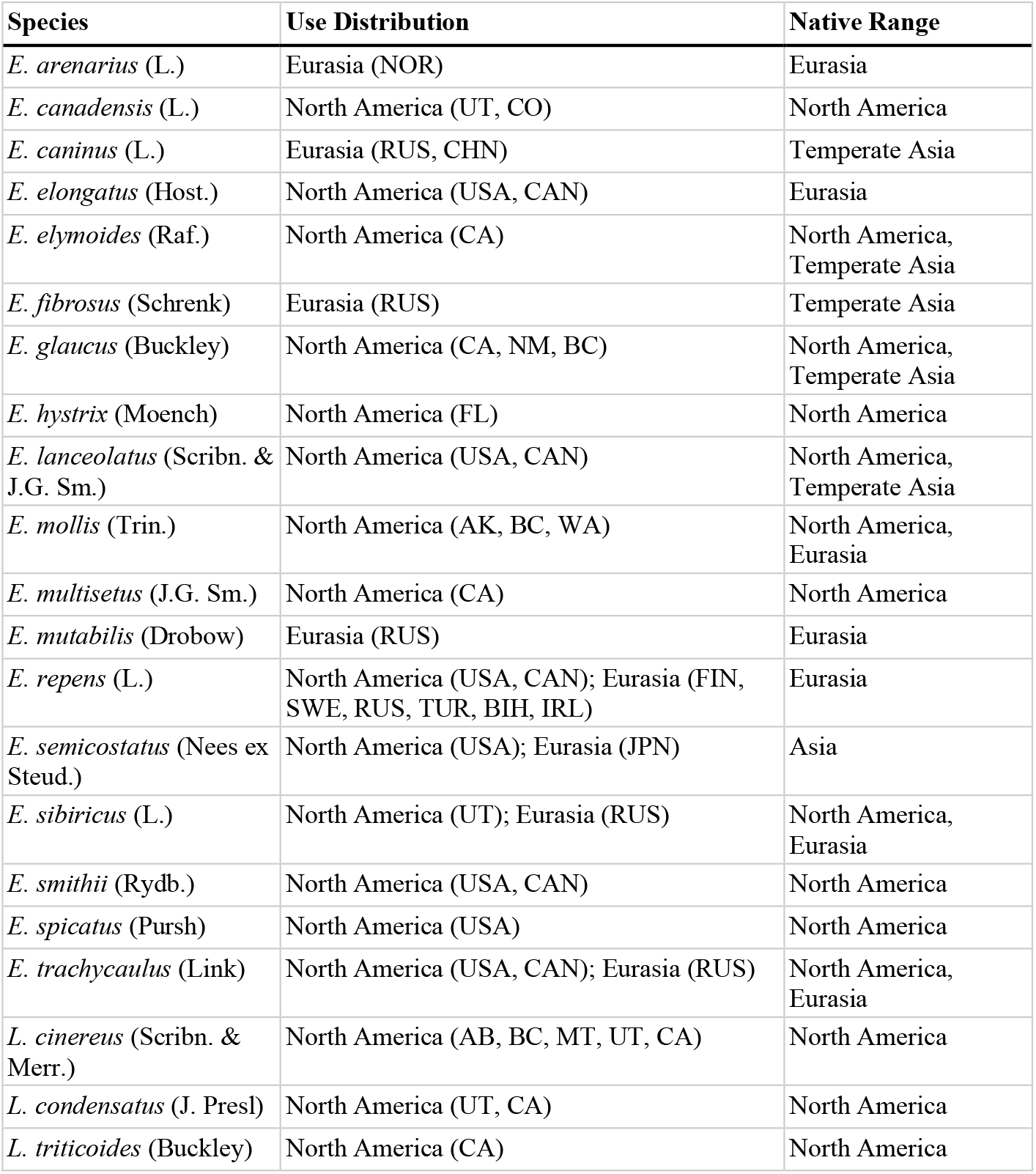
Native ranges and location of ethnobotanical use for 21 *Elymus* species.

**Figure 1.**
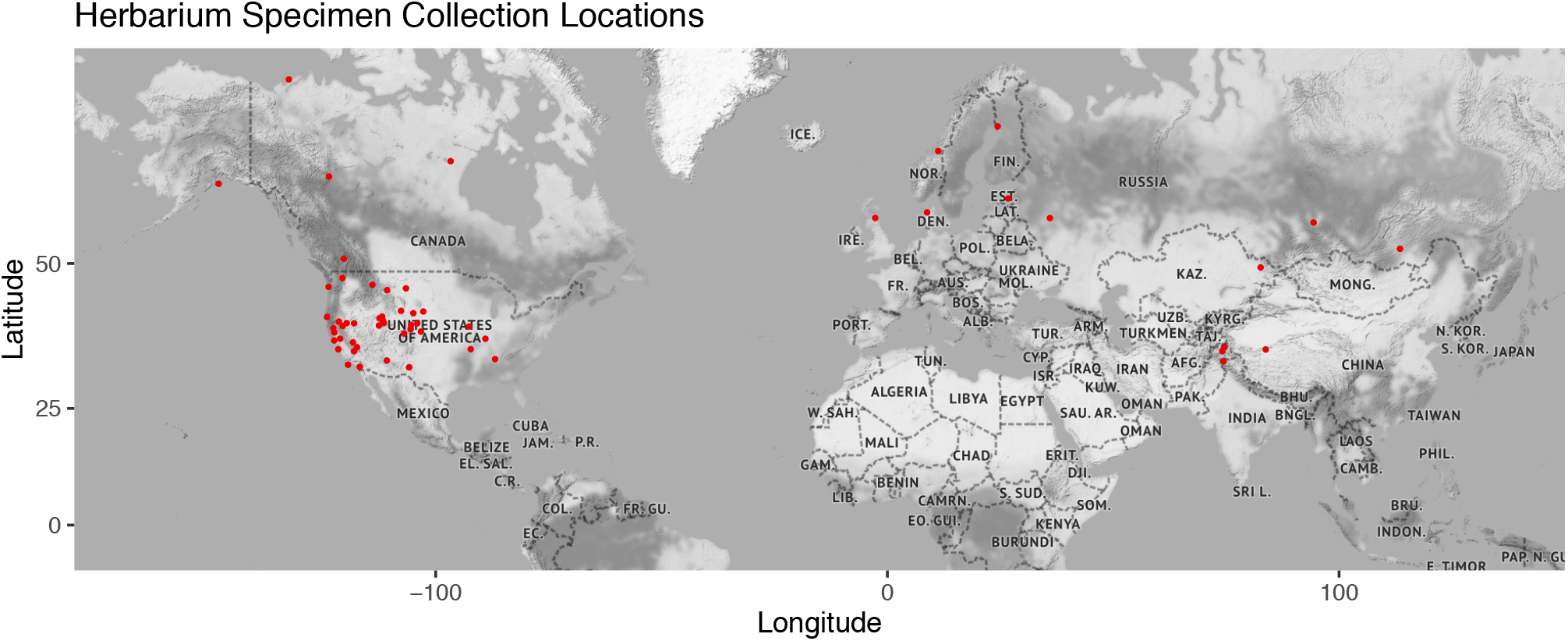
Geographic locations of collection sites for all specimens measured across 21 *Elymus* species. Collection site determined from herbarium specimen label.

To investigate inter- and intra-specific variation in floret area in species used by people we sampled three herbarium specimens per species and harvested eight florets from each specimen, with the exception of *E. semicostatus* Nees ex Steud., for which only two herbarium specimens existed. For every specimen, we recorded the location of collection, accession and collection number, collection date, collector, and latitude and longitude when available (Appendix 1). We removed the glumes to reveal the caryopsis enclosed by the adherent palea and the lemma. We imaged florets in high resolution (iPhone XR, DPI 326) at the Missouri Botanical Garden herbarium and measured area in ImageJ (v. 1.50i). We returned plant material to the fragment packet on the herbarium sheet following imaging. We cropped each image to encompass only the seeds, then converted the image to binary to analyze particles for individual and average floret area (mm^2^). Raw data for floret area is available in Appendix 2. We fit linear models in R (v. 1.0.143, RStudio Team 2015) and SAS (v. 9.4, SAS Institute 2017) to investigate three main questions: 1) does individual floret area differ between species and among replicates within a species, 2) do average floret areas vary with ethnobotanical uses in a given region, and 3) is there an association between average floret area and latitude and longitude? While *Elymus hystrix* Moench was described in the literature as being used medicinally, its specific application (maize seed germination: Table 2) was not consistent with the other species’ medicinal uses. Therefore we removed *E. hystrix* when testing for an effect of medicinal usage on floret size.

**Table 2.**
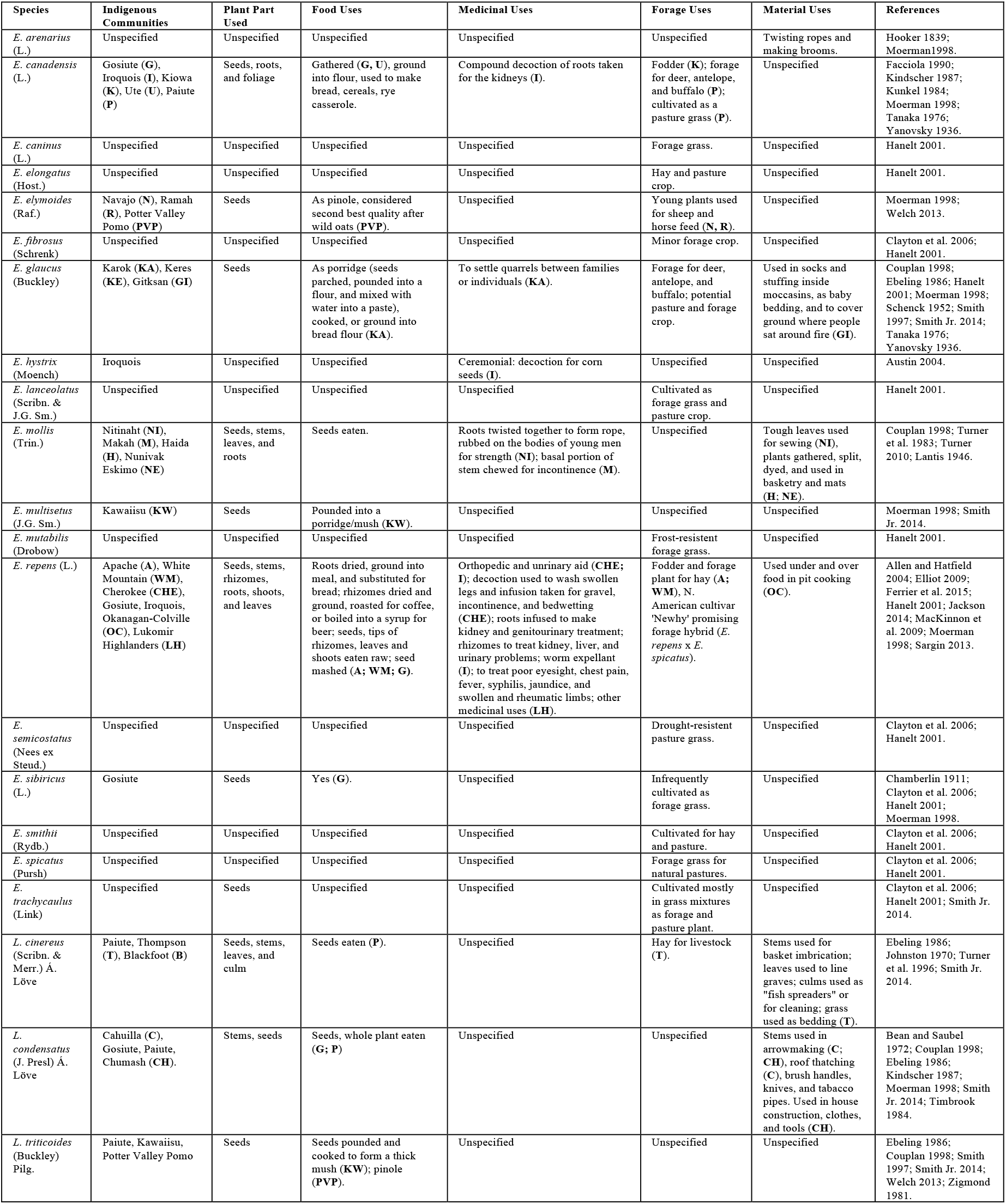
Compilation of documented ethnobotanical records for 21 *Elymus* species. “Unspecified” denotes where an indigenous community, plant part, or ethnobotanical use was not documented for a given species in the literature we consulted.

## Results

### Ethnobotanical analysis of Elymus

Of the ca. 150 known *Elymus* species, we identified 21 taxa that have documented ethnobotanical uses by people in North America and/or Eurasia (Table 2). Fifteen species are used as forage, 12 are used for food, six provide for raw materials for use in the home, and five are used medicinally. We identified at least 25 different indigenous communities that use *Elymus* in some capacity. Five Native American communities use more than one species from the genus: Gosiute (four species), Paiute (four), Kawaiisu (two), Potter Valley Pomo (two), and Iroquois (two). Additionally, eight taxa in our study are used by more than one indigenous group (*E. canadensis, E. elymoides* Raf.*, E. glaucus, E. mollis, E. repens, L. cinereus, L. condensatus*, and *L. triticoides.* Forage uses are mainly as fodder, hay, and pasture grass. Food uses primarily involve the seed, eaten raw (i.e. *E. repens*), as porridge or mash (i.e. *E. glaucus, E. multisetus* J.G. Sm.*, E. repens*), or as ‘pinole,’ a coarse flour made from ground seeds (i.e. *E. elymoides*). Material uses are broad and encompassed many plant parts (culms, leaves, roots, and stems), most frequently as components of houseware (i.e. basketry, broom handles). Medicinal uses are equally diverse, with species being used in decoctions, infusions, and washes.

### Elymus species used for forage

The most common ethnobotanical use of *Elymus* in our study is forage. Fifteen *Elymus* species are used for forage by at least one of seven indigenous communities across western North America (Table 2). Forage uses are primarily for pasture grass, hay for livestock, and fodder for antelope, buffalo deer, horses, and sheep. Seven species are used exclusively as forage (there were no ethnobotanical records of use for food, medicine, or material for the species), whereas eight of the *Elymus* species used for forage are also human edible (Table 2). Many *Elymus* species used as forage have specific environmental tolerances. For example, *E. elongatus* Host. is used as a saline and alkaline tolerant pasture grass in western North America; *E. canadensis* and *E. smithii* Rydb. are used for revegetation and reseeding of disturbed rangelands, prairies, and saline soils of the Great Plains; and *E. lanceolatus* aids soil stabilization in the intermountain region of the United States and Canada (Hanelt 2001). Two additional pasture grass species (*E. mutabilis* Drobow and *E. semicostatus*) are cultivated for their frost and drought resistance, respectively (Hanelt 2001). Finally, *E. elymoides* is edible to sheep and horses early in the season and is used for this purpose by at least two southwestern Native American communities, the Navajo and Ramah (Barkworth 2007; Moerman 1998).

### Elymus species used for human consumption

Ten *Elymus* species are consumed by people in some form, and we identified six indigenous communities that used *Elymus* for this purpose (Table 2). *Elymus* species eaten by humans are: *E. canadensis, E. elymoides, E. glaucus, E. mollis, E. multisetus, E. repens, E. sibiricus* L.*, L. cinerius, L. condensatus*, and *L. triticoides*. For some species there is no comprehensive description of preparation method (i.e. *E. mollis*). Several others illuminate important details on food use; for example, seeds, roots, rhizomes, and leaves of *E. repens* are consumed, either eaten raw, roasted, as a mash, or in a flour. Likewise, seeds of *E. glaucus, E. multisetus*, and *L. triticoides* are parched, ground, and mixed with water to form a type of porridge. Pinole is also a common preparation method for seed, and it is used as a flour in breads (*E. canadensis, E. elymoides, E. glaucus, L. triticoides*), cereals, and casseroles (*E. canadensis*). Notably, *E. elymoides* is considered “second in quality for [pinole]” following wild oats (Welch 2013).

### Elymus species used for medicines and materials

*Elymus* medicinal uses vary widely. Three taxa are used to treat renal and incontinence issues as a diuretic, and two are applied topically to treat swollen limbs (Table 2). *Elymus glaucus* is described by the Karok community as a medicine to help “settle quarrels” between individuals or families (Moerman 1998; Schenk and Gifford 1952). In other medicinal applications, roots and stems are either eaten, applied directly, or developed into infusions and washes. *Elymus hystrix* is described as a “ceremonial medicine” by the Iroquois, and functions as part of a decoction for maize seeds to enhance germination. The treatment is considered to contribute to seed vitality and “protection” prior to planting (Austin 2004; Waugh 1916). Six taxa have material applications (Table 2). Plants are formed into parts of household objects, such as brooms, baskets, arrows, pipes, bedding, brush handles, knives, and mats, among other tools, or into parts of the house, such as in roof thatching. For example, North American Thompson River Indians imbricate stems of *L. cinereus* into baskets (Turner 1996), and *E. arenarius* L. is formed into in ropes and brooms in parts of Eurasia (Hooker 1839). We found that roots, stems, leaves, and culms of *Elymus* are all employed in material ways.

### Floret area measurements

Floret area was measured for 21 *Elymus* species with documented use histories (see above). Floret area varies significantly across *Elymus* species (*F*_20_ = 13.37, *P* < 0.0001), as well as among individuals within species (*F*_21_ = 10.60, *P* < 0.0001). Using species’ means, we fit linear models to assess if average floret area differed by location (i.e. North America vs. Eurasia) and within each of the four ethnobotanical categories (i.e. documented use vs. unspecified) (Table 2). For medicine, floret area does not differ by region (*F*_1_ = 3.63, *P* > 0.05; Figure 2). However, average floret area is significantly greater for species with medicinal uses compared to species without documented medicinal uses in North America (*F*_1_ = 4.75, *P* = 0.03; Figure 2a). In contrast, for food, forage, and material categories, floret area does not differ significantly by region (Food: *F*_1_ = 4.01, *P* > 0.05; Forage: *F*_1_ = 3.93, *P* > 0.05; Material: *F*_1_ = 3.87, *P* > 0.05). Additionally, no differences in floret area are observed when we compare average floret area for species used for food, forage, and material to those without documented usage in each category, (Food: *F*_1_ = 2.33, *P* > 0.05; Forage: *F*_1_ = 1.71, *P* > 0.05; Material; *F*_1_ = 1.24, *P* > 0.05). In summary, average floret area does not differ significantly across geographic regions and among documented ethnobotanical uses, with the exception of species used for medicine in North America. Florets of species used medicinally were larger than florets of species not used medicinally in this region.

**Figure 2:**
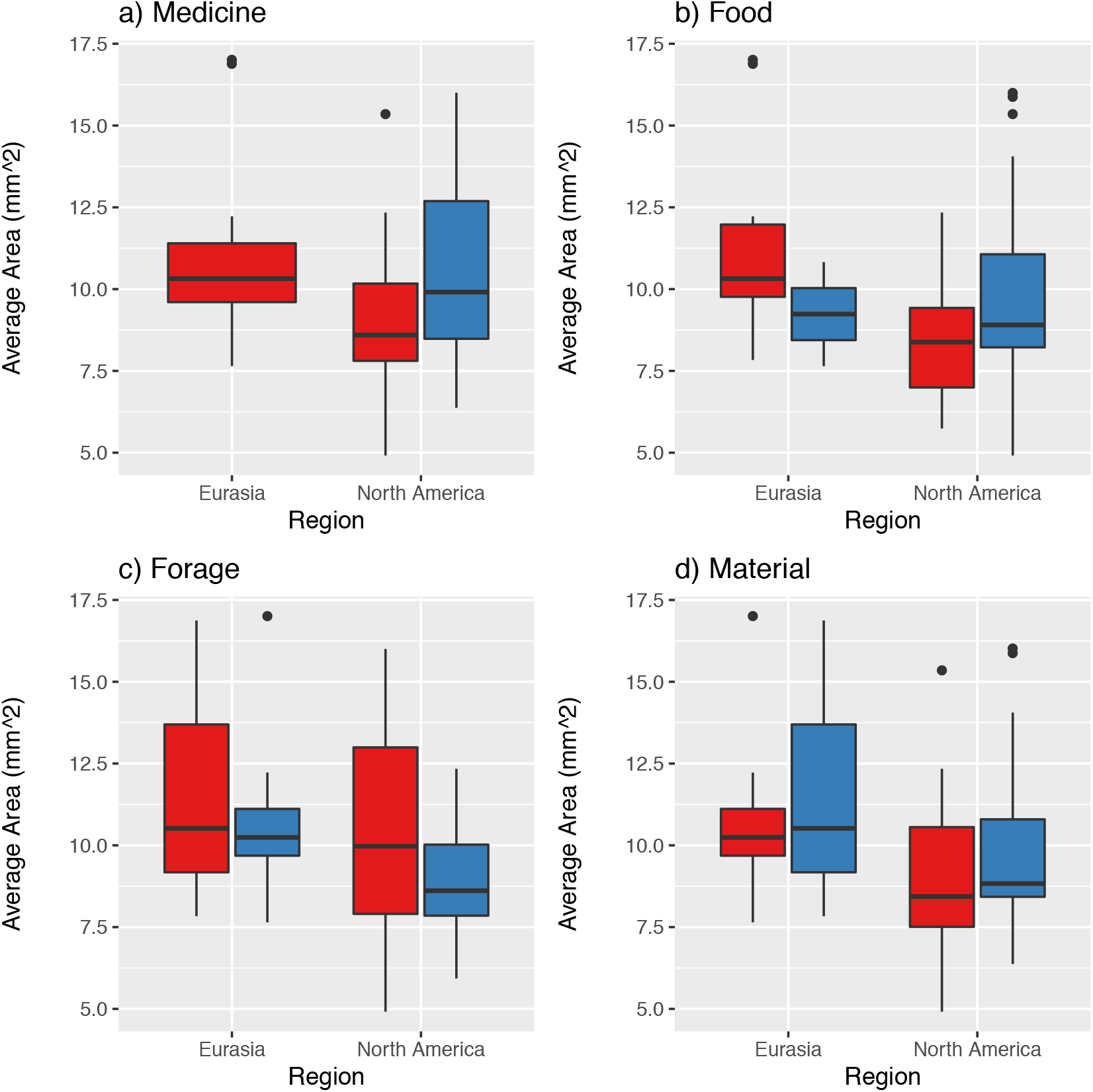
Comparison of average floret area by use (medicine, food, forage, and material) and region (North America, Eurasia). Blue denotes a documented use within that ethnobotanical category. Red denotes no documented use within that ethnobotanical category. Significant differences (*F*_1_ = 4.75, *P* = 0.03) found only for medicinal uses in North America (2a).

To further investigate drivers of variation in average floret area across specimens, we tested for associations between average floret area, latitude, and longitude within region (i.e. North America vs. Eurasia). In North America, average floret area increases from east to west (*t* = −2.41, *P* = 0.02), but is not variable across latitudes (*t* = 1.52, *P* = 0.14). In Eurasia, there is no significant relationship between average floret area and latitude (*t* = 0.20, *P* = 0.85) or longitude (*t* = 0.17, *P* = 0.87); however, this may be an artifact of lower sampling in Eurasia in this study.

## Discussion

Growing concerns about ecological impacts of agricultural systems based on annual plants has turned attention to the potential of perennial, herbaceous species in contemporary food systems. Through their large, persistent root systems, among other traits, perennial, herbaceous plants offer ecological services including reduced erosion and increased absorption of water. However, because of a dearth of herbaceous perennial crops, identifying potential candidates for the ecological intensification of agriculture remains a challenge (Bommarco 2013). Ethnobotanical records play an important role in this process by providing practical information about wild plant use and morphology. *Elymus* is a genus of interest for pre-breeding and domestication processes because of its rich ethnobotanical record, documented edibility, and reproductive morphology. In addition, its history of hybridization suggests that members of the genus may be intercrossed to develop new agricultural cultivars with beneficial trait combinations. While there is no indication that native users of this group selected species with larger floret areas for consumption, forage, or as material, significant variation in floret area exists among and within species. Grain morphology is a valuable target of selection for domestication in perennial grasses, and standing phenotypic variation in this group could serve as a foundation for future breeding initiatives. Moreover, variation in use of *Elymus* species illuminates the potential for broad application of this genus.

### Ethnobotanical analyses as a foundation for agricultural innovation

Ethnobotanical records are a vital source of information on plant diversity, use, distribution, form, and function. In particular, ethnobotanical records can inform agricultural processes by examining how plants have been manipulated or altered for human use (Casas et al. 1996). Further, these studies document which species were chosen for economic and cultural purposes (Ford 2000). Additionally, ethnobotanical resources provide insight on geographic distributions, environmental tolerances, toxicities, preparation methods, and human preferences for certain features (flavors, shapes, textures, colors, etc.) of wild food plants (Casas et al. 1996). These records thereby help identify species with agricultural potential, and provide pertinent information on plant morphology and edibility in an agricultural context (Ciotir et al. 2019; Minnis 2000; Plucknett and Smith 1986).

Our dataset identified 21 species of *Elymus* with known food, forage, medicine, and material uses globally, and attributed these uses to at least 25 different indigenous communities. The most frequent use of *Elymus* is as forage or fodder, further highlighting perennial members of Triticeae as globally important sources of forage grass (Kole 2011). We identified ten species of *Elymus* that are used for food (*E. canadensis, E. elymoides, E. glaucus, E. mollis, E. multisetus, E. repens, E. sibiricus, L. cinerius, L. condensatus* and *L. triticoides*). Of these, the seed is most frequently consumed, though in two instances (e.g. *E. repens* and *L. condensatus*) there are food uses for the entire plant, including the roots and rhizomes (Table 2). The preparation methods for seed are straightforward (i.e. ground and mixed with water as a mash, or finely pounded into flour), suggesting that their edibility is not contingent on rigorous processing. Further, food products prepared with *Elymus* are similar to many modern grain products, and include bread, flour, and cereal. We suggest further investigation of these ten species for their potential contribution to the ecological intensification of agriculture. Lastly, the documentation of medicinal and material uses suggests that *Elymus* taxa are multifunctional, and perhaps the whole plant can be employed post-production or at the end of their lifespan (i.e. as hay for livestock or in thatching).

A previous ethnobotanical study of annual and perennial wild grass genera substantiated their importance as a food source for Native American communities, including species from *Oryzopsis, Sporobolus*, and *Panicum* and highlighted their potential to elucidate cereal domestication processes (Doebley 1984). Similarly, ethnobotanical studies of other wild foods have resulted in recommendations for their agricultural improvement, such as in grain chenopods (Partap and Kapoor 1985). Other studies suggest improved collections of wild plants to encourage their cultivation, such as in wild onion (*Allium*) (Bye 1985). Thus, in addition to identifying potential crops, ethnobotanical studies can result in a variety of suggestions for pre-breeding and domestication efforts in wild food plants.

*Elymus* is a cosmopolitan genus, and the 21 species in this study with documented ethnobotanical uses have widely distributed native ranges, occurring across temperate North America and Eurasia. While we identified documented uses for *Elymus* in several Eurasian countries (Table 1), the depth of ethnobotanical information about *Elymus* species used in North America was much greater. This disparity could be accredited to the fact that we primarily used resources at the Missouri Botanical Garden library, thereby biasing the details of our study to North America and to resources in English. As such, there are other globally-distributed *Elymus* species that could have been used in an ethnobotanical capacity and that may also have potential for use in pre-breeding and domestication programs. For example, wild relatives of sunflower (Heliantheae) with larger ranges may have environmental tolerances and other traits useful to breeding initiatives (Kantar et al. 2015).

### Variation in floret traits of Elymus species

Seed traits (floret traits in *Elymus*) are an important feature of wild and domesticated plants that may bear some indication of other agronomically and ecologically important features. For example, wild taxa with larger seeds can have larger seedlings, faster rates of germination, higher recruitment success, and greater reproductive output, though trade-offs in seed size and seed number exist for some species (Giles 1990; Jakobsson and Eriksson 2000). Similarly, during grain domestication, selection favors species and individuals with larger seeds, resulting in greater seedling vigor, root and shoot biomass, and yield, though the correlation of seed size to plant size at maturity is weaker (Milla and Matesanz 2017; Preece et al. 2015; Rees and Venable 2007; Stougaard and Xue 2004). Further, it has been found that the progenitors of cereal crops have larger seeds than other wild grasses that have never undergone domestication (Preece et al. 2015). Given this information, we hypothesized that *Elymus* species with larger floret areas would perhaps be more frequently used as a food source and be more desirable for human consumption/domestication purposes. For the *Elymus* taxa examined in this study, we found significant differences in floret area between species and among replicates of a species. This suggests that there is substantial natural variation in floret area within *Elymus*, an economically and agriculturally important trait with potential for selection and evolution through the pre-breeding process. From a conservation perspective, these data underscore the importance of a dynamic *in situ* and *ex situ* conservation management that targets multiple species within a genus, and diverse populations in different geographic locations (e.g., Khoury et al. 2019).

We observed a significant relationship between floret area and medicinal use in North American *Elymus* species. We cannot ensure that the specimens measured accurately reflect the plants used by indigenous communities in the last three centuries. However, some studies exploring seeds traits of medicinal plants assess seed size in relation to oil content (i.e. *Moringa*, Mani et al. 2007; *Pentaclethra*, Asoegwu et al. 2006). The medicinal uses of vegetative plant parts (i.e. roots) of *Elymus* exist, yet seeds were rarely described in a medicinal context (Table 2); therefore, the benefit of a larger floret area for medicinal applications should be further investigated. For example, what properties of *Elymus* grains matter in medicinal applications (oils, carbohydrates)? Are the grains ground, infused, or eaten directly in a medicinal context? Future work could use voucher specimens from ethnobotanical studies to track the relationship between medicinal use and floret area. Further, this study and others like it emphasize the importance of plant use histories in conservation management, as different cultural communities have unique and varied uses for the same species or closely related species; or, they have used different species for similar purposes (e.g., Albuquerque et al. 2009)

Despite the wide variation in floret size within and among *Elymus* species, we did not observe a significant relationship between average floret area, region, and documented ethnobotanical use for the remaining three categories examined here (food, forage, and material). It is conceivable that use of *Elymus* species for forage and material would not necessarily lead to changes in seed size, as the primary structures being used (e.g., stems, leaves) may have been the targets of selection. Data for these components of the plant were not collected in this study; consequently, we are unable to assess whether or not forage and material uses led to changes in these traits. Regarding food uses, archaeological analyses of taxa previously used for food have demonstrated differences in seed size and other traits over time (Langlie et al. 2014; Mueller 2017), providing evidence for selection and domestication. The lack of association between floret size and use of *Elymus* for food in our dataset indicates that floret area was not associated with utilization or consumption by indigenous communities. Detailed analyses of other traits, including inflorescence size, plant height, historical abundance, may provide insights into selection during at their time of use. Nonetheless, nearly all of the collections sampled in this study were from the 20th century, and floret areas may have varied more significantly at the time and place of use. Further, comparative analyses of the *Elymus* species with documented use histories with other *Elymus* species for which no use history is known, might shed light on how floret traits in *Elymus* species have changed through their interaction with humans. Floret and grain traits remain important for *de novo* domestication in grasses and should be examined more extensively in *Elymus* as well as other taxa of interest.

## Conclusions

Morphological and genetic variation in cultivated plants, their wild progenitors, and other wild species provides the foundation for plant domestication and breeding efforts. In response to concerns about long-term sustainability of our current agricultural system, attention is focusing in part on *de novo* domestication of wild species (Ciotir et al. 2016; Ciotir et al. 2019; Kole 2011). As such, *Elymus* and many other genera of herbaceous perennials, merits increased attention to its research, development, and conservation. These efforts include improving the availability of *Elymus* germplasm in biorepositories globally in conjunction with expanding the collection of ethnobotanical histories throughout the genus. Additionally, we suggest more comprehensive morphological and molecular studies of taxa with documented food uses to more precisely identify promising candidates for agriculture. Similarly, we see value in *in-situ* conservation for genetically, phenotypically, and culturally valuable populations (i.e. at sites of indigenous use), as well as in-ground plantings to assess survivability in a controlled environment. Ultimately, a variety of *Elymus* species show promise for the ecological intensification of agriculture.

## Acknowledgements

This research was supported by The Perennial Agriculture Project in conjunction with the Malone Family Land Preservation Foundation and The Land Institute, the Missouri Botanical Garden, and Saint Louis University. The authors are grateful to Dr. James Solomon and Dr. Gerrit Davidse for their help in the Missouri Botanical Garden herbarium. We thank Dr. Laura Klein and Zachary Harris for valuable comments on the manuscript.

## Author Contributions

AJM, BM, CC, ESF, and MJR conceived of the study. BM, CC, and ESF collected the data. ESF and MJR performed data analyses. AJM, BM, and ESF took the lead in writing the manuscript and all authors contributed to manuscript editing.

## Appendix Captions

### Appendix 1 caption

Herbarium specimen information from which florets were harvested for area measurements. * = Specific latitudes and longitudes were not available at time of collection, so coordinates were estimated in Google Earth for general analyses in R based off of detailed geographic information provided on the specimen.

### Appendix 2 caption

Individual floret area measurements for 21 *Elymus* species and replicates.

## References

Albuquerque, U.P., Sousa Araújo, T.A., Ramos, M.A., Teixeira do Nascimento, V., Lucena, R.F.P., Monteiro, J.M., Alencar, N.L., E. Lima Araújo. 2009. How Ethnobotany Can Aid Biodiversity Conservation: Reflections on Investigations in the Semi-Arid Region of NE Brazil. Biodiversity and Conservation 18(1): 127–50. https://doi.org/10.1007/s10531-008-9463-8.

Asoegwu, S., Ohanyere, S., Kanu, O.P., C.N. Iwueke. 2006. Physical Properties of African Oil Bean Seed (*Pentaclethra macrophylla*). Agricultural Engineering International: the CIGR Ejournal. Manuscript FP 05006. Vol. VIII.

Aubry, C., Shoal, R., V. Erickson. 2005. Grass cultivars: their origins, development, and use on national forests and grasslands in the Pacific Northwest. Pendleton (OR): USDA Forest Service, Umatilla National Forest. Available at: https://www.fs.fed.us/wildflowers/Native_Plant_Materials/documents/cultivars_maindoc_040405_appendices.pdf

Austin, D. F. 2004. Florida Ethnobotany. Boca Raton, FL: CRC.

Awika, J.M. 2011. Major Cereal Grains Production and Use around the World. Advances in Cereal Science.

Barkworth, M.E. 2007. Triticeae. In: Flora of North America Magnoliophyta: Commelinidae (in part): Poaceae, Part 1: North of Mexico. Vol 24, pp. 239–374. Oxford, Oxford University Press.

Bean, L. J., K.S. Saubel. 1972. Temalpakh: Cahuilla Indian knowledge and usage of plants. Banning: Malki Museum.

Bommarco, R., Kleijn, D., S.G. Potts. 2013. Review: Ecological intensification: harnessing ecosystem services for food security. Trends in Ecology & Evolution, 28(4): 230–238. https://doi:10.1016/j.tree.2012.10.012.

Bye, R. A. 1985. Botanical perspectives of ethnobotany of the Greater Southwest. Economic Botany: 39(4), 375–386. doi: 10.1007/bf02858744

Casas, A., del Carmen Vázquez, M., Viveros, J.L., J. Caballero. 1996. Plant Management Among the Nahua and the Mixtec in the Balsas River Basin, Mexico: An Ethnobotanical Approach to the Study of Plant Domestication. Human Ecology, 24(4): 455–478.

Cassman, K.G. 1999. Ecological intensification of cereal production systems: Yield potential, soil quality, and precision agriculture. Proceedings National Academy of Sciences, 96(11): 5952–5959. https://doi.org/10.1073/pnas.96.11.5952.

Chamberlin, L.V. 1911. The ethnobotany of the Gosiute Indians. Academy of Natural Sciences, 61(1): 24–99.

Chen, S., G., Zhu. 2006. Poaceae (Tribe Triticeae). Wu Z.Y. and Raven P.H. (eds.). Flora of China, Vol. 22, Poaceae. 386–663. Science Press, Beijing, and Missouri Botanical Garden Press, St. Louis.

Ciotir, C., Applequist, W., Crews, T.E., Cristea, N., DeHaan, L.R., Frawley, E., Herron, S., Magill, R., Miller, J., Roskov, Y., Schlautman, B., Solomon, J., Townesmith, A., Van Tassel, D., Zarucchi, J., A.J. Miller. 2019. Building a botanical foundation for perennial agriculture: Global inventory of wild, perennial herbaceous Fabaceae species. Plants, People, Planet, 00: 1–12. https://doi.org/10.1002/ppp3.37

Clayton, W.D., Vorontsova, M.S., Harman, K.T., H. Williamson. 2006 onwards. GrassBase - The Online World Grass Flora. http://www.kew.org/data/grasses-db.html

Couplan, F. 1998. The Encyclopedia of Edible Plants of North America. Keats Publishing.

Cox, T.S., Bender, M., Picone, C., Van Tassel, D.L., Holland, J.B., Brummer, E.C., Zoeller, B.E., Paterson, A.H., W. Jackson. 2002. Breeding perennial grain crops. Critical Reviews in Plant Sciences, 21(2): 59–91.

Cox, T.S., Glover, J.D., Van Tassel, D.L., Cox, C.M., L.R. DeHaan. 2006. Prospects for developing perennial grain crops. BioScience, 56(8): 649–659.

Cox, T.S. 2009. Chapter 1: Crop domestication and the first plant breeders. Pp 1–26 in S. Ceccarelli; E.P. Guimar; E. Weltizien (eds.), Plant breeding and farmer participation. U.N. Food and Agriculture Organization.

Cox, S., Nabukalu, P., Paterson, A. Kong, W., S. Nakasagga. 2018. Development of Perennial Grain Sorghum. Sustainability, 10(1): 172–180. https://doi.org/10.3390/su10010172.

Doebley, J. F. 1984. “Seeds” of wild grasses: A major food of Southwestern Indians. Economic Botany, 38(1): 52–64. doi:10.1007/bf02904415

Doré, T., Makowski, D., Malézieux, E., Munier-Jolain, N., Tchamitchian, M., P. Tittonell. 2011. Facing up to the paradigm of ecological intensification in agronomy: Revisiting methods, concepts and knowledge. European Journal of Agronomy, 34: 197–210. https://doi:10.1016/j.eja.2011.02.006.

DeHaan, L.R., Christians, M., Crain, J. Poland. 2018. Development and Evolution of an Intermediate Wheatgrass Domestication Program. Sustainability, 10(5): 1499. https://doi.org/10.3390/su10051499

DeHaan, L. R., D. L. Van Tassel, J. A. Anderson, S. R. Asselin, R. Barnes, G. J. Baute, D. J. Cattani, S. W. Culman, K. M. Dorn, B. S. Hulke, M. Kantar, S. Larson, M. D. Marks, A. J. Miller, J. Poland, D. A. Ravetta, E. Rude, M. R. Ryan, D. Wyse, X. Zhang. 2016. A Pipeline Strategy for Grain Crop Domestication. Crop Science, 56: 917–930. doi: 10.2135/cropsci2015.06.0356

DeHaan, L.R., D.L. Van Tassel. 2014. Useful insights from evolutionary biology for developing perennial grain crops. American Journal of Botany, 101(10): 1801–1819.

DeHaan, L. R., Van Tassel D. L., T.S. Cox. 2010. Missing domesticated plant forms: can artificial selection fill the gap? Evolutionary Applications, 3(5-6): 434–452. doi: 10.1111/j.1752-4571.2010.00132.x

Ebeling, W. 1986. Handbook of Indian foods and fibers of arid America. Berkeley u.a.: Univ. of Calif. Pr.

Elliot, B.A. 2009. Handbook of edible and poisonous plants of western North America. Elliot Environmental Consulting, LLC.

Facciola. S. 1990. Cornucopia: A Source Book of Edible Plants. California: Kampong Publications.

Fatih, A.M.B. 1983. Analysis of the breeding potential of wheat-Agropyron and wheat-Elymus derivatives. Hereditus, 98: 287–295.

Ferrier, J., Saciragic, L., Trakic, S., Chen, E.C.H., Gendron, R.L., Cuerrier, A., Balick, M.J., Redzic, S., Alikadic, E., J.T. Arnason. 2015. An ethnobotany of the Lukomir Highlanders of Bosnia & Herzegovina. Journal of Ethnobiology and Ethnomedicine, 11: 81.

Food and Agriculture Organization of the United Nations (FAO). 2009. A Global Treaty for Food Security and Sustainable Agriculture, International Treaty on plant genetic resources for food and agriculture. FAO, Rome: Electronic Publishing Policy and Support Branch Communication Division.

Food and Agriculture Organization of the United Nations (FAO). 2019. The State of the World’s Biodiversity for Food and Agriculture, J. Bélanger & D. Pilling (eds.). FAO Commission on Genetic Resources for Food and Agriculture Assessments. Rome. (http://www.fao.org/3/CA3129EN/CA3129EN.pdf) Licence: CC BY-NC-SA 3.0 IGO.

Ford, R.I. 2000. Agriculture: An Introduction. in Minnis, P.E. (ed.) Ethnobotany: A Reader. University of Oklahoma Press, Norman. Pp. 243.

Füleky, G. (ed.) 2009. Cultivated Plants, Primarily As Food Sources. Eolss Publishers Co. Ltd. Oxford, United Kingdom.

Gegas, V.C., Nazari, A., Griffiths, S., Simmonds, J., Fish, L., Orford, S., Sayers, L., Doonan, J.H., J.W. Snape. 2010. A genetic framework for grain size and shape variation in wheat. Plant Cell, 22(4): 1046–55. doi: 10.1105/tpc.110.074153.

Giles, B.E. 1990. The effects of variation in seed size on growth and reproduction in the wild barley Hordeum vulgare ssp. Spontaneum. Heredity 64: 239–250.

Gregg, B.R., G.L. Billups. 2010. Seed Conditioning, Volume 3: Crop Seed Conditioning. CRC Press. IBSN: 1439846715.

Glémin, S., T. Bataillon. 2009. A comparative view of the evolution of grasses under domestication. New Phytologist, 183: 273–290.

Glover, J. D., J. P. Reganold, L. W. Bell, J. Borevitz, E. C. Brummer, E. S. Buckler, C. M. Cox, et al. 2010. “Increased Food and Ecosystem Security via Perennial Grains.” Science 328 (5986): 1638–39. https://doi.org/10.1126/science.1188761.

Hanelt, P. 2001. Mansfeld’s Encyclopedia of Agricultural and Horticultural Crops, vol. 5. New York, Berlin: Springer.

Harlan J., de Wet J., E. Price. 1973. Comparative evolution of cereals. Evolution, 27: 311–325. doi:10.1111/j.1558-5646.1973.tb00676.x.

Harlan, J.R. 1992. Crops and Man, 2nd Edition. American Society of Agronomy and Crop Science Society of America, Madison, WI.

Hayes, R.C., Wang, S., Newell, M.T., Turner, K., Larsen, J., Gazza, L., Anderson, J.A., Bell, L.W., Cattani, D.J., Frels, K., Galassi, E., Morgounov, A.I., Revell, C.K., Thapa, D.B., Sacks, E.J., Sameri, M., Wade, L.J., Westerbergh, A., Shamanin, V., Amanov, A., G.D. Li. 2018. The performance of early-generation perennial winter cereals at 21 sites across four continents. Sustainability, 10(4):1124–1152. https://doi.org/10.3390/su10041124.

Helfgott, D.E, R.J. Mason-Gamer. 2004. The evolution of North America *Elymus* (Triticeae, Poaceae) allotetraploids: evidence from phosphoenolpyruvate carboxylase gene sequences. Systematic Botany, 29(4): 850–861.

Hooker, W.D. 1839. Notes on Norway; or a brief journal of a tour made to the Northern parts of Norway, in the summer of MDCCCXXXVI.

Huang, G., Qin, S., Zhang, S., Cai, X.,Wu, S., Dao, J., Zhang, J., Huang, L., Harnpichitvitaya, D., Wade, L., F. Hu. 2018. Performance, economics and potential impact of perennial rice PR23 relative to annual rice cultivars at multiple locations in Yunnan Province of China. Sustainability, 10(4): 1086–1104. https://doi.org/10.3390/su10041086.

Jakobsson, A., O. Eriksson. 2000. A comparative study of seed number, seed size, seedling size and recruitment in grassland plants. Oikos, 88(3): 494–502. doi:10.1034/j.1600-0706.2000.880304.x

Johnston, A. 1970. Blackfoot Indian utilization of the flora of the northwestern Great Plains. Economic Botany, 24(3): 301–324.

Kantar, M. B., Sosa, C.C., Khoury, C.K., Castañeda-Álvarez, N.P., Achicanoy, H.A., Bernau, V. Kane, N.C., Marek, L., Seiler, G., L. H. Rieseberg. 2015. Ecogeography and Utility to Plant Breeding of the Crop Wild Relatives of Sunflower (Helianthus Annuus L.). Frontiers in Plant Science 6: 841. https://doi.org/10.3389/fpls.2015.00841.

Kellogg, E.A. 2015. The Families and Genera of Vascular Plants: Flowering Plants, Monocots, Poaceae. Switzerland, Springer International Publishing.

Khoury, C. K., Amariles, D., Soto, J.S., Diaz, M.V., Sotelo, S., Sosa, C.C., Ramírez-Villegas, J., et al. 2019. Comprehensiveness of Conservation of Useful Wild Plants: An Operational Indicator for Biodiversity and Sustainable Development Targets. Ecological Indicators 98: 420–29. https://doi.org/10.1016/j.ecolind.2018.11.016

Khoury, C.K., Bjorkman, A.D., Dempewoldf, H., Ramirez-Villegas, J., Guarino, L., Jarvis, A., Rieseberg, L.H., P.C. Struik. 2014. Increasing homogeneity in global food supplies. Proceedings of the National Academy of Sciences, 111(11): 4001–4006. doi:10.1073/pnas.1313490111.

Kole, C. 2011. Wild crop relatives: genomic and breeding resources; cereals. Berlin Heidelberg, Springer-Verlag.

Kunkel, G. 1984. Plants for human consumption: An Annotated Checklist of the Edible Phanerogams and Ferns. Koeltz Scientific Books, Koenigstein.

Langlie, B.S., Mueller, N.G., Spengler, R.N., G.J. Fritz. 2014. Agricultural Origins From the Ground Up: Archaeological Approaches to Plant Domestication. American Journal of Botany 101(10): 1601–1617.

Lantis, M. 1946. The social culture of the Nunivak Eskimo. Transactions of the American Philosophical Society, 35(3): 153–323.

Lloyd-Reilley, J. 2010. Plant guide for Canada wildrye (*Elymus canadensis*). USDA-Natural Resources Conservation Service, E. “Kika” de la Garza Plant Materials Center. Kingsville, TX. Available at: https://plants.usda.gov/plantguide/pdf/pg_elca4.pdf [accessed June 21, 2019].

Lu, B.R. 1993. Biosystematic investigations of asiatic wheatgrasses-*Elymus* L. (*Triticeae: Poaceae*). The Swedish University of Agricultural Sciences.

MacKinnon et al. 2009. Edible & medicinal plants of Canada. Lone Pine Publishing.

Mani, S., Jaya, S., R. Vadivambal. 2007. Optimization of Solvent Extraction of Moringa *(Moringa Oleifera)* Seed Kernel Oil Using Response Surface Methodology. Food and Bioproducts Processing 85(4): 328–335. https://doi.org/10.1205/fbp07075

Mueller, N.G. 2017. Veget Hist Archaeobot 26: 313. https://doi.org/10.1007/s00334-016-0592-9

Milla, R., S. Matesanz. 2017. Growing larger with domestication: a matter of physiology, morphology or allocation? Plant Biology, 19(3), 475–483. doi:10.1111/plb.12545

Miller, A.J., Novy, A., Glover, J., Kellogg, E.A., Maul, J.E., Raven, P., P. Wyse Jackson. 2015. Expanding the role of botanical gardens in the future of food. Nature Plants, 1(15078): doi: 10.1038/nplants.2015.78.

Millet, E. 1986. Relationships Between Grain Weight and the Size of Floret Cavity in the Wheat Spike. Annals of Botany, 58(3): 417–423. http://www.jstor.org/stable/42757682

Millet, E., M.J. Pinthus. 1984. The association between grain volume and grain weight in wheat. Cereal Science, 2(1): 31–35. https://doi.org/10.1016/S0733-5210(84)80005-3

Minnis, P.E. 2000. Ethnobotany: A Reader. University of Oklahoma Press, Norman.

Moerman, D. E. 1998. Native American ethnobotany. Timber Press, Portland.

National Geographic Society (NGS). 2008. Edible: an illustrated guide to the world’s food plants. National Geographic Society, Washington D.C.

Okamoto, Y. Kajimura, T. Ikeda, T.M., S. Takumi. 2012. Evidence from principal component analysis for improvement of grain shape and spikelet morphology related traits after hexaploid wheat speciation. Genes Genet Syst, 87(5): 299–310.

Olsen, K.M., J.F. Wendel. 2013a. Crop plants as models for understanding plan adaptation and diversification. Frontiers in Plant Science, 1(4): 290. doi: 10.3389/fpls.2013.00290.

Olsen, K.M., J.F. Wendel. 2013b. A bountiful harvest: genomic insights into crop domestication phenotypes. Annual Reviews in Plant Biology, 64: 47–70. DOI: 10.1146/annurev-arplant-050312-120048.

Partap, T., P. Kapoor. 1985. The Himalayan Grain Chenopods: Distribution and Ethnobotany. Agriculture, Ecosystems, and Environment, 14(3-4): 185–199. https://doi.org/10.1016/0167-8809(85)90035-0

Plucknett, D.L., N.J.H. Smith. 1986. International prospects for cooperation in crop research. Economic Botany, 40(3): 298–309. doi:10.1007/bf02858987

Preece, C., Livarda, A., Wallace, M., Martin, G., Charles, M., Christin, P.A., Jones, G., Rees, M., C.P. Osborne. 2015. Were Fertile Crescent crop progenitors higher yielding than other wild species that were never domesticated? New Phytologist, 207(3): 905–913.

Rees M., D.L. Venable. 2007. Why do big plants make big seeds? Journal of Ecology 95: 926–936.

RStudio Team. 2015. RStudio: Integrated Development for R. RStudio, Inc., Boston, MA. http://www.rstudio.com/.

Sargin, S.A., Akçicek, E., S. Selvi. 2013. An ethnobotanical study of the medicinal plants used by the local people of Alasehir (Manisa) in Turkey. Journal of Ethnopharmacology, 150(3): 860–874.

SAS Institute, Base SAS 9.4 Procedures Guide: Statistical Procedures, Fifth Edition. SAS Institute, 2017.

Schenck, S.M., E. W. Gifford. 1952. Karok Ethnobotany. Anthropological Records, 13(6): 377–392.

Smith, H. 1997. Ethnobotany of the Gitksan Indians of British Columbia. Canadian Museum of Civilization.

Smith Jr., J. P. 2014. Field guide to grasses of California. University of California Press.

St. John, L., Jensen, K., Ogle, D.G., D. Tilley. 2010. Plant guide for RS Hybrid wheatgrass (*Elymus hoffmannii*). USDA-Natural Resources Conservation Service, Aberdeen, ID Plant Materials Center and USDA-ARS Forage and Range Laboratory, Logan, UT. Available at: https://plants.usda.gov/plantguide/pdf/pg_elho3.pdf

Stougaard, R.N., Q.W. Xue. 2004. Spring wheat seed size and seeding rate effects on yield loss due to wild oat (*Avena fatua*) interference. Weed Science, 52: 133–141.

Tanaka, T. 1976. Tanaka’s cyclopaedia of edible plants of the world. Keigaku Publishing.

Tilley, D., Ogle, D., L. St. John. 2011. Plant guide for slender wheatgrass (*Elymus trachycaulus* ssp. *trachycaulus*). USDA-Natural Resources Conservation Service, Idaho Plant Materials Center. Aberdeen, ID. https://plants.usda.gov/plantguide/pdf/pg_eltr7.pdf

Timbrook, J. 1984. Chumash Ethnobotany: A preliminary report. Journal of Ethnobiology, 4(2): 141–169.

Tittonell, P. 2014. Ecological intensification of agriculture — sustainable by nature. Current Opinion in Environmental Sustainability, 8: 53–61. https://doi.org/10.1016/j.cosust.2014.08.006

Turner, N.J. 2010. Plants of Haida Gwaii. Winlaw, B.C.: Sono Nis Press.

Turner, N.J., Thomas, J., Carlson, B.F., R.T. Ogilvie. 1983. Ethnobotany of the Nitinaht Indians of Vancouver Island. Victoria: Ministry of Provincial Secretary and Government Services, Province of British Columbia.

Turner, N.J., Thompson, L.C., Thompson, M.T., A.Z. York. 1996. Thompson ethnobotany: knowledge and usage of plants by the Thompson Indians of British Columbia. Victoria, B.C: Royal British Columbia Museum.

Wang, S.L., Qi, L.L., Chen, P.D., Liu, D.J., Friebe, B., B.S. Gill. 1999. Molecular cytogenetic identification of wheat-Elymus tsukushiense introgression lines. Euphytica, 107: 217–224.

Warren, J. 2015. The nature of crops: how we came to eat the plants we do. Wallingford: CABI. 1780645082.

Waugh, F.W. 1916. Iroquois Foods and Food Preparation. University Press of the Pacific. ISBN 1410207765.

Welch, J.R. 2013. Sprouting valley historical ethnobotany of the northern pomo from Potter Valley, California. Denton, TX: Society of Ethnobiology.

Yanovsky, E. 1936. Food plants of the North American Indians. Publication no. 237. U.S. Dept of Agriculture.

Zigmond, M.L. 1981. Kawaiisu Ethnobotany. Salt Lake City: Univ. of Utah Press.

Zohary, D., Hopf, M. E. Weiss (eds.). 2012. Domestication of Plants in the Old World. 4th edition. New York: Oxford University Press.

