## Appendix 1 for "An ethnobotanical study of the genus *Elymus*"

| **Species** | **Replicate** | **Location** | **Accession #** | **Collection #** | **Latitude** | **Longitude** | **Date Collected** | **Collector** | **Locality Notes** |
| --- | --- | --- | --- | --- | --- | --- | --- | --- | --- |
| *E. arenarius* | 1 | Europe (Norway) | 2235195 | NA | 63.47 | 11.27 | 5/8/96 | J.E. Elsley | Shore of Trondheimsfjord. |
| *E. canadensis* | 1 | North America (Montana) | 907705 | 4965 | 46.33333333 | -106.5833333 | 7/6/78 | E.E. Terrell | N/A |
| *E. caninus* | 1 | Asia (Russia) | 3294987 | NA | 56.035483* | 35.97126* | Year 1991 | A. K. Skvortsov | European province of Moscow, district Volokolamsk. |
| *L. cinereus* | 1 | North America (California) | 6732874 | 2791 | 36.58333333 | -117.4166667 | 7/17/84 | P.M. Peterson, C.R. Annable | N/A |
| *L. condensatus* | 1 | North America (Nevada) | 1215729 | 2723 | 40.75277778 | -119.7027778 | 5/26/41 | A.A. Beetle | N/A |
| *E. elongatus* | 1 | North America (Utah) | 4813335 | 15763 | 40.77472222 | -111.4688889 | 7/12/99 | S.A. Thompson, B.L. Isaac | Summit County, UT. |
| *E. elymoides* | 1 | North America (California) | 6603991 | 11108 | 39.89666667 | -122.6466667 | 6/18/04 | L. Ahart | Tehama County, CA. |
| *E. fibrosus* | 1 | Asia (Russia) | 2371341 | 4948a | NA | NA | 1914 | NA | N/A |
| *E. glaucus* | 1 | North America (Canada) | 744524 | 76821 | 62.45416667 | -96.69027778 | 6/15/08 | J. Macoun | N/A |
| *E. hystrix* | 1 | North America (Alabama) | 3442275 | 68728 | 34.45 | -86.85 | 6/9/82 | R. Kral | N/A |
| *E. lanceolatus* | 1 | North America (Colorado) | 1123131 | 1121 | NA | NA | 8/1/34 | A. Nelson, R. Nelson | Sand dunes of North Park, CO. |
| *E. mollis* | 1 | North America (Washington) | 5061104 | 370 | 46.56666667 | -123.7166667 | 7/3/91 | Arnot | N/A |
| *E. multisetus* | 1 | North America (California) | 4153962 | 3323 | 36.232621 | -121.549455 | 6/1/01 | A.D. E. Elmer | Tassajara Hot Springs, Monterey CA. |
| *E. mutabilis* | 1 | Asia (Russia) | 5200096 | 167 | 55.48333333 | 94.36666667 | 7/16/88 | E. Antipova | Siberia, Krasnoyarsk Region. |
| *E. repens* | 1 | North America (Colorado) | 4822734 | 13849 | 39.73694444 | -105.5397222 | 7/2/05 | R.M. King, R. M. Garvey | N/A |
| *E. semicostasus* | 1 | Asia (Kashmir) | 1047872 | 10554 | 34.047434* | 74.394091* | 8/31/28 | R.R. Stewart | Kashmir near Tangmarg. |
| *E. sibiricus* | 1 | North America (Canada) | 1803394 | 11953 | 60.83333333 | -123.6166667 | 8/3/61 | W.J. Cody, K.W. Spicer | N/A |
| *E. smithii* | 1 | North America (Missouri) | 4305774 | 5663 | 40.19027778 | -92.60638889 | 6/18/70 | M. L. Conrad | N/A |
| *E. spicatus* | 1 | North America (Washington) | 1082729 | 10 782 | 47.88333333 | -120.6333333 | NA | J.W. Thompson | N/A |
| *E. trachycaulus* | 1 | North America (Colorado) | 6482682 | 14364 | 40.73833333 | -104.0680556 | 5/29/06 | R.M. King, R. M. Garvey | N/A |
| *L. triticoides* | 1 | North America (California) | 2964728 | 3572 | 33.03333333 | -116.8 | 5/24/03 | L.R. Abrams | N/A |
| *E. arenarius* | 2 | Europe (Scotland) | 1826000 | 17 | 56.055803* | -2.705749* | 8/8/56 | P.S. Green | East links near North Berwick, East Lothian. Sand dune. |
| *E. canadensis* | 2 | North America (Colorado) | 6721689 | 11993 | 40.46055556 | -105.4030556 | 9/13/92 | C.R. Annable | N/A |
| *E. caninus* | 2 | Asia (Khazakstan) | 5036661 | NA | 49.43337* | 82.628396* | 8/18/70 | J.A. Kotukhou | Kalbinsky Range, Sibinskie lakes. Kazakh Altai Mountain. |
| *L. cinereus* | 2 | North America (Montana) | 2377325 | 826 | 46.05* | -110.7666667* | 8/23/21 | W.N. Suksdorf | Suksdorf Gulch, 9 miles NW of Wilsall. |
| *L. condensatus* | 2 | North America (California) | 4287116 | 4807 | 33.41666667* | -119.4* | 10/20/92 | J. Ricketson, T. Schmidt | Los Angeles County, Santa Catalina Conservancy. |
| *E. elongatus* | 2 | North America (Nebraska) | 2619376 | 1605 | 42.681256* | -102.733859* | 7/8/73 | S.P. Churchill | 2 miles east of Hay Springs 100 meters south |
| *E. elymoides* | 2 | North America (California) | 6215215 | 10550 | 35.88222222 | -118.0744444 | 6/27/03 | S. Boyd | Southern Sierra Nevada. |
| *E. fibrosus* | 2 | Europe (Finland) | 1710472 | 1827 | 65.806027* | 24.412727* | 1826 | E. Hayren | Kaakamo, Finland. |
| *E. glaucus* | 2 | North America (California) | 950210 | 4530 | 41.78333333* | -124.05* | 6/5/28 | J.W. Thompson | Douglas Park, Del Norte County CA. |
| *E. hystrix* | 2 | North America (Arkansas) | 1267030 | 23533 | 36.251827* | -92.23986* | 7/14/42 | P.O. Ellis | Above Norfork Dam, Baxter County AR. |
| *E. lanceolatus* | 2 | North America (Nevada) | 6647035 | 16604 | 40.73333333 | -118.0666667 | 6/21/13 | A. Tiehm | N/A |
| *E. mollis* | 2 | North America (Alaska) | 5394597 | 5937 | 60.05 | -148.05 | 7/25/48 | W.J. Eyerdam | Port San Juan, Evans Island. |
| *E. multisetus* | 2 | North America (California) | 983472 | 1523 | 38.1* | -121.15* | 5/26/30 | E.E. Stanford | Harney Lane, San Joaquin County CA. |
| *E. mutabilis* | 2 | Asia (Russia) | 5164252 | NA | NA | NA | 7/11/71 | A.C. Pebymknh | N/A |
| *E. repens* | 2 | North America (Arizona) | 5743483 | NA | 34.2 | -110.8 | 7/31/03 | F. E. Northam | N/A |
| *E. semicostasus* | 2 | Asia (Pakistan) | 6048420 | 13757 | 35.91666667 | 74.16666667 | 9/25/95 | B. Dickore | N/A |
| *E. sibiricus* | 2 | Asia (Russia) | 4954205 | NA | 52.05* | 113.4833333* | 7/16/72 | NA | Chita, East towards Baykal Lake. |
| *E. smithii* | 2 | North America (Montana) | 6313576 | 211 | 46.872162* | -114.013155* | 7/7/56 | E. E. Addor | Missoula Valley, MT. California St. and Arlington St. intersection. |
| *E. spicatus* | 2 | North America (New Mexico) | 4372477 | 69-323 | 32.946274* | -105.878011* | 8/19/69 | N. C. Henderson | 10 miles east of Cloudcroft, NM. Along US 82 in Otero County. |
| *E. trachycaulus* | 2 | North America (Utah) | 5398024 | 12099 | 41.200438* | -111.693023* | 8/12/55 | F.J. Herman | Weber County, southwest of Huntsville, Rocky summit of ridge |
| *L. triticoides* | 2 | North America (California) | 6614044 | 10209 | 37.78333333 | -122.4666667 | 7/28/90 | P. Rubtzoff, N. Rubtzoff | Presidio San Fran County, SW of Marine Hospital, N side of Lobos Creek. |
| *E. arenarius* | 3 | Europe (Denmark) | 3922335 | 793 | 56.731518* | 8.792154* | 6/25/75 | E. Boel, B. Ollgaard | Island of Mors, SE of Hojris. |
| *E. canadensis* | 3 | North America (Utah) | 1710635 | 8136 | 41.85* | -111.85* | 7/18/50 | A. H. Holmgren | Cache County, UT. |
| *E. caninus* | 3 | Europe (Estonia) | 2358366 | NA | 58.377443* | 26.720348* | 2/4/05 | NA | Tartu, Estonia. |
| *L. cinereus* | 3 | North America (British Columbia) | 2620785 | 1158 | 50.68333333* | -120.3333333* | 6/28/72 | A. G. Jones | North Thompson River Railroad, Kamloops. |
| *L. condensatus* | 3 | North America (California) | 1215206 | 6582 | 41* | -121.4166667* | 5/25/40 | C.L. Hitchcock | Shasta County, C. 2 miles west of Fall River Mills. |
| *E. elongatus* | 3 | North America (Wyoming) | 1740678 | 4956 | 42.776841* | -107.674274* | 7/6/49 | C.L. Porter | 10 miles east of Sand Draw Oil Field, 40 miles SE of Riverton. Fremont County, WY. |
| *E. elymoides* | 3 | North America (California) | 802045 | 12357 | 39.18333333* | -122.4333333* | 6/6/16 | A.A. Heller | 3 miles west of Leesville, Colsua County, CA. |
| *E. fibrosus* | 3 | Europe (Finland) | 2745303 | NA | 66.142388* | 25.077544* | 7/29/78 | Y. Ulvinen | Tervola, N shore of Kemijoki River. |
| *E. glaucus* | 3 | North America (Colorado) | 1641429 | 2602 | 39* | -107.0333333* | 8/4/49 | G.B. Van Shaack | Valley below Schofield Pass in Gunnison National Forest. |
| *E. hystrix* | 3 | North America (Illinois) | 1882658 | 1499 | 38.089548* | -88.994522* | 7/1/67 | T.S. Elias | 1 mile east of Sesser on Route 183. |
| *E. lanceolatus* | 3 | North America (Utah) | 6070617 | 18-Jul | 40.344874* | -112.560422* | 6/21/07 | A. Kelsey, S. Sedivec | In Stansbury Mountains, 1/4 mile east of Johnson Pass. |
| *E. mollis* | 3 | North America (Canada) | 5942098 | 13155 | 69.73333333 | -132.5 | 8/1/63 | W.J. Cody | N/A |
| *E. multisetus* | 3 | North America (California) | 4153943 | 8329 | 37.45* | -118.3166667* | 5/26/06 | J.G. Smith | White Mountains. Southern Bell mine, Mono County, CA. |
| *E. mutabilis* | 3 | Asia (Pakistan) | 5668224 | 98-274 | 36.58333333 | 74.58333333 | 7/12/98 | E. Eberhardt | Batura Valley. |
| *E. repens* | 3 | North America (Utah) | 3139297 | 4797 | 41.58333333* | -111.9166667* | 7/11/83 | S. L. Hatch | 3 miles south of Wellsville, Eldon Cooper Ranch. |
| *E. sibiricus* | 3 | Asia (China) | 4373360 | 1908 | 36.20309* | 83.749671* | 6/26/88 | S.G. Wu, H. Ohba, W.H. Wu, Y. Fei | Xinjiang, Kulun Mountain. |
| *E. smithii* | 3 | North America (Colorado) | 1125999 | 1345 | 39.327103* | -103.274499 | 7/13/37 | M. Ownbey | 2 miles north of Arriba, Lincoln County, CO. |
| *E. spicatus* | 3 | North America (Utah) | 3318363 | 13,136 | 41.557875* | -112.428132* | 6/27/60 | A. Holmgren, W.S. Boyle, B. Palmer | 12 miles south of Howell in Box Elder County, UT. |
| *E. trachycaulus* | 3 | North America (Wyoming) | 5446962 | 8358 | 42.40837* | -104.944324* | 7/2/01 | A. Nelson | Cassa, Laramie County, WY. |
| *L. triticoides* | 3 | North America (California) | 2986139 | 49522 | 40.28333333* | -120.5666667* | 6/28/73 | J.T. Howell, G.H. True | Lassen County, CA. Elysian Valley, west of Janesville. |

**Appendix 1**: Herbarium specimen information from which florets were harvested for area measurements. * = Specific latitudes and longitudes were not available at time of collection, so coordinates were estimated in Google Earth for general analyses in R based off of detailed geographic information provided on the specimen.
