## Appendix 2 for "An ethnobotanical study of the genus *Elymus*"

| **Species** | **Replicate** | **Floret Area** |
| --- | --- | --- |
| *Elymus arenarius* | 1 | 14.241 |
| *Elymus arenarius* | 1 | 14.995 |
| *Elymus arenarius* | 1 | 15.036 |
| *Elymus arenarius* | 1 | 15.192 |
| *Elymus arenarius* | 1 | 15.235 |
| *Elymus arenarius* | 1 | 16.894 |
| *Elymus arenarius* | 1 | 17.116 |
| *Elymus arenarius* | 1 | 26.273 |
| *Elymus canadensis* | 1 | 7.859 |
| *Elymus canadensis* | 1 | 9.016 |
| *Elymus canadensis* | 1 | 9.724 |
| *Elymus canadensis* | 1 | 10.564 |
| *Elymus caninus* | 1 | 8.016 |
| *Elymus caninus* | 1 | 8.432 |
| *Elymus caninus* | 1 | 8.573 |
| *Elymus caninus* | 1 | 9.556 |
| *Elymus caninus* | 1 | 9.774 |
| *Elymus caninus* | 1 | 11.736 |
| *Elymus caninus* | 1 | 12.265 |
| *Elymus elongatus* | 1 | 6.637 |
| *Elymus elongatus* | 1 | 7.933 |
| *Elymus elongatus* | 1 | 8.197 |
| *Elymus elongatus* | 1 | 10.22 |
| *Elymus elongatus* | 1 | 10.738 |
| *Elymus elongatus* | 1 | 11.769 |
| *Elymus elongatus* | 1 | 12.573 |
| *Elymus elongatus* | 1 | 14.862 |
| *Elymus elymoides* | 1 | 9.043 |
| *Elymus elymoides* | 1 | 10.36 |
| *Elymus elymoides* | 1 | 10.538 |
| *Elymus elymoides* | 1 | 11.422 |
| *Elymus elymoides* | 1 | 11.887 |
| *Elymus elymoides* | 1 | 12.85 |
| *Elymus elymoides* | 1 | 13.787 |
| *Elymus fibrosus* | 1 | 7.751 |
| *Elymus fibrosus* | 1 | 7.95 |
| *Elymus fibrosus* | 1 | 9.135 |
| *Elymus fibrosus* | 1 | 9.522 |
| *Elymus fibrosus* | 1 | 10.17 |
| *Elymus fibrosus* | 1 | 10.225 |
| *Elymus fibrosus* | 1 | 11.975 |
| *Elymus fibrosus* | 1 | 12.059 |
| *Elymus glaucus* | 1 | 8.87 |
| *Elymus glaucus* | 1 | 9.525 |
| *Elymus glaucus* | 1 | 9.633 |
| *Elymus glaucus* | 1 | 10.297 |
| *Elymus glaucus* | 1 | 10.343 |
| *Elymus glaucus* | 1 | 10.963 |
| *Elymus glaucus* | 1 | 11.577 |
| *Elymus glaucus* | 1 | 12.987 |
| *Elymus hystrix* | 1 | 9.967 |
| *Elymus hystrix* | 1 | 10.608 |
| *Elymus hystrix* | 1 | 11.271 |
| *Elymus hystrix* | 1 | 11.442 |
| *Elymus hystrix* | 1 | 11.813 |
| *Elymus hystrix* | 1 | 13.564 |
| *Elymus hystrix* | 1 | 14.838 |
| *Elymus lanceolatus* | 1 | 5.987 |
| *Elymus lanceolatus* | 1 | 6.508 |
| *Elymus lanceolatus* | 1 | 8.76 |
| *Elymus lanceolatus* | 1 | 9.862 |
| *Elymus mollis* | 1 | 9.961 |
| *Elymus mollis* | 1 | 12.218 |
| *Elymus mollis* | 1 | 13.558 |
| *Elymus mollis* | 1 | 14.864 |
| *Elymus mollis* | 1 | 15.114 |
| *Elymus mollis* | 1 | 15.149 |
| *Elymus mollis* | 1 | 22.996 |
| *Elymus mollis* | 1 | 23.152 |
| *Elymus multisetus* | 1 | 10.537 |
| *Elymus multisetus* | 1 | 11.834 |
| *Elymus multisetus* | 1 | 12.511 |
| *Elymus multisetus* | 1 | 14.604 |
| *Elymus multisetus* | 1 | 15.015 |
| *Elymus multisetus* | 1 | 17.546 |
| *Elymus multisetus* | 1 | 17.928 |
| *Elymus multisetus* | 1 | 22.742 |
| *Elymus mutabilis* | 1 | 9.452 |
| *Elymus mutabilis* | 1 | 10.791 |
| *Elymus mutabilis* | 1 | 10.856 |
| *Elymus mutabilis* | 1 | 11.703 |
| *Elymus mutabilis* | 1 | 12.112 |
| *Elymus mutabilis* | 1 | 13.211 |
| *Elymus mutabilis* | 1 | 13.676 |
| *Elymus mutabilis* | 1 | 13.982 |
| *Elymus repens* | 1 | 4.268 |
| *Elymus repens* | 1 | 4.907 |
| *Elymus repens* | 1 | 6.129 |
| *Elymus repens* | 1 | 6.178 |
| *Elymus repens* | 1 | 6.881 |
| *Elymus repens* | 1 | 7.852 |
| *Elymus repens* | 1 | 8.394 |
| *Elymus semicostatus* | 1 | 11.045 |
| *Elymus semicostatus* | 1 | 12.007 |
| *Elymus semicostatus* | 1 | 15.26 |
| *Elymus semicostatus* | 1 | 18.614 |
| *Elymus semicostatus* | 1 | 20.033 |
| *Elymus semicostatus* | 1 | 20.068 |
| *Elymus semicostatus* | 1 | 21.934 |
| *Elymus sibiricus* | 1 | 6.371 |
| *Elymus sibiricus* | 1 | 6.676 |
| *Elymus sibiricus* | 1 | 7.596 |
| *Elymus sibiricus* | 1 | 11.846 |
| *Elymus sibiricus* | 1 | 12.35 |
| *Elymus smithii* | 1 | 5.995 |
| *Elymus smithii* | 1 | 6.105 |
| *Elymus smithii* | 1 | 6.901 |
| *Elymus smithii* | 1 | 7.085 |
| *Elymus smithii* | 1 | 7.963 |
| *Elymus smithii* | 1 | 9.582 |
| *Elymus smithii* | 1 | 9.841 |
| *Elymus smithii* | 1 | 14.014 |
| *Elymus spicatus* | 1 | 6.988 |
| *Elymus spicatus* | 1 | 9.634 |
| *Elymus spicatus* | 1 | 9.705 |
| *Elymus spicatus* | 1 | 10.967 |
| *Elymus spicatus* | 1 | 11.083 |
| *Elymus spicatus* | 1 | 11.824 |
| *Elymus spicatus* | 1 | 11.905 |
| *Elymus spicatus* | 1 | 14.758 |
| *Elymus trachycaulus* | 1 | 3.26 |
| *Elymus trachycaulus* | 1 | 5.096 |
| *Elymus trachycaulus* | 1 | 5.492 |
| *Elymus trachycaulus* | 1 | 5.561 |
| *Elymus trachycaulus* | 1 | 5.757 |
| *Elymus trachycaulus* | 1 | 6.464 |
| *Elymus trachycaulus* | 1 | 7.824 |
| *Elymus trachycaulus* | 1 | 7.984 |
| *Leymus cinereus* | 1 | 6.649 |
| *Leymus cinereus* | 1 | 7.251 |
| *Leymus cinereus* | 1 | 7.331 |
| *Leymus cinereus* | 1 | 7.401 |
| *Leymus cinereus* | 1 | 8.387 |
| *Leymus cinereus* | 1 | 9.861 |
| *Leymus cinereus* | 1 | 11.107 |
| *Leymus cinereus* | 1 | 12.759 |
| *Leymus condensatus* | 1 | 7.413 |
| *Leymus condensatus* | 1 | 9.791 |
| *Leymus condensatus* | 1 | 11.807 |
| *Leymus condensatus* | 1 | 12.041 |
| *Leymus condensatus* | 1 | 12.226 |
| *Leymus condensatus* | 1 | 12.415 |
| *Leymus triticoides* | 1 | 6.119 |
| *Leymus triticoides* | 1 | 6.465 |
| *Leymus triticoides* | 1 | 7.087 |
| *Leymus triticoides* | 1 | 7.236 |
| *Leymus triticoides* | 1 | 7.271 |
| *Leymus triticoides* | 1 | 8.06 |
| *Leymus triticoides* | 1 | 8.697 |
| *Leymus triticoides* | 1 | 10.286 |
| *Elymus arenarius* | 2 | 8.256 |
| *Elymus arenarius* | 2 | 8.848 |
| *Elymus arenarius* | 2 | 10.078 |
| *Elymus arenarius* | 2 | 10.306 |
| *Elymus arenarius* | 2 | 10.854 |
| *Elymus arenarius* | 2 | 11.488 |
| *Elymus arenarius* | 2 | 12.071 |
| *Elymus arenarius* | 2 | 12.25 |
| *Elymus canadensis* | 2 | 9.231 |
| *Elymus canadensis* | 2 | 10.481 |
| *Elymus canadensis* | 2 | 10.935 |
| *Elymus canadensis* | 2 | 11.156 |
| *Elymus canadensis* | 2 | 11.91 |
| *Elymus canadensis* | 2 | 13.222 |
| *Elymus canadensis* | 2 | 13.631 |
| *Elymus canadensis* | 2 | 13.857 |
| *Elymus caninus* | 2 | 9.918 |
| *Elymus caninus* | 2 | 10.974 |
| *Elymus caninus* | 2 | 11.772 |
| *Elymus caninus* | 2 | 12.256 |
| *Elymus caninus* | 2 | 12.344 |
| *Elymus caninus* | 2 | 12.38 |
| *Elymus caninus* | 2 | 12.823 |
| *Elymus caninus* | 2 | 15.313 |
| *Elymus elongatus* | 2 | 4.725 |
| *Elymus elongatus* | 2 | 5.308 |
| *Elymus elongatus* | 2 | 6.17 |
| *Elymus elongatus* | 2 | 6.474 |
| *Elymus elongatus* | 2 | 6.766 |
| *Elymus elongatus* | 2 | 8.153 |
| *Elymus elongatus* | 2 | 9.025 |
| *Elymus elymoides* | 2 | 7.572 |
| *Elymus elymoides* | 2 | 8.356 |
| *Elymus elymoides* | 2 | 9.885 |
| *Elymus elymoides* | 2 | 10.738 |
| *Elymus elymoides* | 2 | 10.739 |
| *Elymus elymoides* | 2 | 13.543 |
| *Elymus elymoides* | 2 | 14.378 |
| *Elymus fibrosus* | 2 | 8.295 |
| *Elymus fibrosus* | 2 | 8.332 |
| *Elymus fibrosus* | 2 | 9.504 |
| *Elymus fibrosus* | 2 | 10.214 |
| *Elymus fibrosus* | 2 | 10.863 |
| *Elymus glaucus* | 2 | 7.045 |
| *Elymus glaucus* | 2 | 7.388 |
| *Elymus glaucus* | 2 | 8.399 |
| *Elymus glaucus* | 2 | 8.565 |
| *Elymus glaucus* | 2 | 8.732 |
| *Elymus glaucus* | 2 | 9.478 |
| *Elymus glaucus* | 2 | 9.518 |
| *Elymus glaucus* | 2 | 9.772 |
| *Elymus hystrix* | 2 | 4.99 |
| *Elymus hystrix* | 2 | 5.023 |
| *Elymus hystrix* | 2 | 5.401 |
| *Elymus hystrix* | 2 | 8.16 |
| *Elymus hystrix* | 2 | 8.79 |
| *Elymus lanceolatus* | 2 | 4.466 |
| *Elymus lanceolatus* | 2 | 4.985 |
| *Elymus lanceolatus* | 2 | 6.242 |
| *Elymus lanceolatus* | 2 | 7.409 |
| *Elymus lanceolatus* | 2 | 11.249 |
| *Elymus mollis* | 2 | 8.191 |
| *Elymus mollis* | 2 | 14.349 |
| *Elymus mollis* | 2 | 19.736 |
| *Elymus mollis* | 2 | 21.731 |
| *Elymus multisetus* | 2 | 7.924 |
| *Elymus multisetus* | 2 | 9.255 |
| *Elymus multisetus* | 2 | 9.483 |
| *Elymus multisetus* | 2 | 10.75 |
| *Elymus multisetus* | 2 | 12.452 |
| *Elymus mutabilis* | 2 | 7.52 |
| *Elymus mutabilis* | 2 | 9.153 |
| *Elymus mutabilis* | 2 | 9.698 |
| *Elymus mutabilis* | 2 | 10.26 |
| *Elymus mutabilis* | 2 | 10.689 |
| *Elymus mutabilis* | 2 | 11.301 |
| *Elymus mutabilis* | 2 | 12.532 |
| *Elymus repens* | 2 | 6.768 |
| *Elymus repens* | 2 | 7.013 |
| *Elymus repens* | 2 | 7.017 |
| *Elymus repens* | 2 | 7.3 |
| *Elymus repens* | 2 | 8.29 |
| *Elymus repens* | 2 | 8.533 |
| *Elymus repens* | 2 | 12.369 |
| *Elymus repens* | 2 | 12.794 |
| *Elymus semicostatus* | 2 | 7.729 |
| *Elymus semicostatus* | 2 | 8.874 |
| *Elymus semicostatus* | 2 | 10.067 |
| *Elymus semicostatus* | 2 | 10.186 |
| *Elymus semicostatus* | 2 | 10.534 |
| *Elymus semicostatus* | 2 | 10.621 |
| *Elymus semicostatus* | 2 | 12.063 |
| *Elymus semicostatus* | 2 | 13.751 |
| *Elymus sibiricus* | 2 | 8.329 |
| *Elymus sibiricus* | 2 | 9.598 |
| *Elymus sibiricus* | 2 | 9.65 |
| *Elymus sibiricus* | 2 | 10.314 |
| *Elymus sibiricus* | 2 | 10.489 |
| *Elymus sibiricus* | 2 | 11.503 |
| *Elymus sibiricus* | 2 | 12.846 |
| *Elymus sibiricus* | 2 | 13.885 |
| *Elymus smithii* | 2 | 7.544 |
| *Elymus smithii* | 2 | 8.056 |
| *Elymus smithii* | 2 | 8.25 |
| *Elymus smithii* | 2 | 8.625 |
| *Elymus smithii* | 2 | 11.178 |
| *Elymus smithii* | 2 | 14.403 |
| *Elymus spicatus* | 2 | 11.053 |
| *Elymus spicatus* | 2 | 11.53 |
| *Elymus spicatus* | 2 | 11.746 |
| *Elymus spicatus* | 2 | 12.063 |
| *Elymus spicatus* | 2 | 12.556 |
| *Elymus spicatus* | 2 | 13.693 |
| *Elymus spicatus* | 2 | 13.737 |
| *Elymus trachycaulus* | 2 | 5.783 |
| *Elymus trachycaulus* | 2 | 6.881 |
| *Elymus trachycaulus* | 2 | 7.504 |
| *Elymus trachycaulus* | 2 | 7.91 |
| *Elymus trachycaulus* | 2 | 8.808 |
| *Elymus trachycaulus* | 2 | 9.674 |
| *Elymus trachycaulus* | 2 | 14.132 |
| *Leymus cinereus* | 2 | 5.359 |
| *Leymus cinereus* | 2 | 5.813 |
| *Leymus cinereus* | 2 | 7.741 |
| *Leymus cinereus* | 2 | 8.222 |
| *Leymus cinereus* | 2 | 8.475 |
| *Leymus cinereus* | 2 | 9.42 |
| *Leymus cinereus* | 2 | 10.333 |
| *Leymus cinereus* | 2 | 15.273 |
| *Leymus condensatus* | 2 | 5.564 |
| *Leymus condensatus* | 2 | 6.177 |
| *Leymus condensatus* | 2 | 6.681 |
| *Leymus condensatus* | 2 | 9.698 |
| *Leymus condensatus* | 2 | 9.931 |
| *Leymus condensatus* | 2 | 10.115 |
| *Leymus condensatus* | 2 | 10.246 |
| *Leymus condensatus* | 2 | 11.038 |
| *Leymus triticoides* | 2 | 6.4 |
| *Leymus triticoides* | 2 | 6.489 |
| *Leymus triticoides* | 2 | 7.377 |
| *Leymus triticoides* | 2 | 8.22 |
| *Leymus triticoides* | 2 | 8.36 |
| *Leymus triticoides* | 2 | 9.028 |
| *Leymus triticoides* | 2 | 9.683 |
| *Leymus triticoides* | 2 | 9.7 |
| *Elymus arenarius* | 3 | 6.117 |
| *Elymus arenarius* | 3 | 7.435 |
| *Elymus arenarius* | 3 | 8.146 |
| *Elymus arenarius* | 3 | 9.635 |
| *Elymus canadensis* | 3 | 8.099 |
| *Elymus canadensis* | 3 | 10.048 |
| *Elymus canadensis* | 3 | 13.887 |
| *Elymus canadensis* | 3 | 14.354 |
| *Elymus canadensis* | 3 | 14.779 |
| *Elymus caninus* | 3 | 6.832 |
| *Elymus caninus* | 3 | 7.679 |
| *Elymus caninus* | 3 | 10.362 |
| *Elymus caninus* | 3 | 10.737 |
| *Elymus caninus* | 3 | 10.863 |
| *Elymus caninus* | 3 | 11.1 |
| *Elymus caninus* | 3 | 11.863 |
| *Elymus caninus* | 3 | 13.115 |
| *Elymus elongatus* | 3 | 6.083 |
| *Elymus elongatus* | 3 | 6.759 |
| *Elymus elongatus* | 3 | 6.773 |
| *Elymus elongatus* | 3 | 7.37 |
| *Elymus elongatus* | 3 | 7.488 |
| *Elymus elongatus* | 3 | 7.847 |
| *Elymus elongatus* | 3 | 7.896 |
| *Elymus elongatus* | 3 | 8.7 |
| *Elymus elymoides* | 3 | 6.532 |
| *Elymus elymoides* | 3 | 6.677 |
| *Elymus elymoides* | 3 | 7.536 |
| *Elymus elymoides* | 3 | 8.44 |
| *Elymus glaucus* | 3 | 5.55 |
| *Elymus glaucus* | 3 | 7.319 |
| *Elymus glaucus* | 3 | 7.582 |
| *Elymus glaucus* | 3 | 7.713 |
| *Elymus glaucus* | 3 | 7.726 |
| *Elymus glaucus* | 3 | 7.903 |
| *Elymus glaucus* | 3 | 9.735 |
| *Elymus glaucus* | 3 | 11.226 |
| *Elymus hystrix* | 3 | 4.374 |
| *Elymus hystrix* | 3 | 4.449 |
| *Elymus hystrix* | 3 | 5.036 |
| *Elymus hystrix* | 3 | 5.096 |
| *Elymus hystrix* | 3 | 5.715 |
| *Elymus hystrix* | 3 | 9.762 |
| *Elymus lanceolatus* | 3 | 5.732 |
| *Elymus lanceolatus* | 3 | 6.124 |
| *Elymus lanceolatus* | 3 | 8.048 |
| *Elymus lanceolatus* | 3 | 8.276 |
| *Elymus lanceolatus* | 3 | 9.22 |
| *Elymus lanceolatus* | 3 | 9.37 |
| *Elymus lanceolatus* | 3 | 10.291 |
| *Elymus lanceolatus* | 3 | 10.396 |
| *Elymus mollis* | 3 | 12.757 |
| *Elymus mollis* | 3 | 12.786 |
| *Elymus mollis* | 3 | 13.147 |
| *Elymus mollis* | 3 | 13.161 |
| *Elymus mollis* | 3 | 15.573 |
| *Elymus mollis* | 3 | 16.917 |
| *Elymus multisetus* | 3 | 3.979 |
| *Elymus multisetus* | 3 | 4.395 |
| *Elymus multisetus* | 3 | 4.642 |
| *Elymus multisetus* | 3 | 4.909 |
| *Elymus multisetus* | 3 | 4.933 |
| *Elymus multisetus* | 3 | 5.344 |
| *Elymus multisetus* | 3 | 5.431 |
| *Elymus multisetus* | 3 | 5.698 |
| *Elymus mutabilis* | 3 | 5.373 |
| *Elymus mutabilis* | 3 | 7.325 |
| *Elymus mutabilis* | 3 | 7.7 |
| *Elymus mutabilis* | 3 | 7.893 |
| *Elymus mutabilis* | 3 | 9.566 |
| *Elymus mutabilis* | 3 | 9.773 |
| *Elymus mutabilis* | 3 | 10.854 |
| *Elymus repens* | 3 | 5.627 |
| *Elymus repens* | 3 | 5.867 |
| *Elymus repens* | 3 | 7.325 |
| *Elymus repens* | 3 | 7.346 |
| *Elymus repens* | 3 | 8.852 |
| *Elymus repens* | 3 | 8.969 |
| *Elymus repens* | 3 | 9.169 |
| *Elymus repens* | 3 | 9.76 |
| *Elymus sibiricus* | 3 | 5.405 |
| *Elymus sibiricus* | 3 | 6.717 |
| *Elymus sibiricus* | 3 | 7.112 |
| *Elymus sibiricus* | 3 | 7.508 |
| *Elymus sibiricus* | 3 | 7.569 |
| *Elymus sibiricus* | 3 | 9.417 |
| *Elymus sibiricus* | 3 | 9.796 |
| *Elymus smithii* | 3 | 6.266 |
| *Elymus smithii* | 3 | 7.278 |
| *Elymus smithii* | 3 | 8.258 |
| *Elymus smithii* | 3 | 8.545 |
| *Elymus smithii* | 3 | 8.774 |
| *Elymus smithii* | 3 | 10.868 |
| *Elymus spicatus* | 3 | 4.905 |
| *Elymus spicatus* | 3 | 5.343 |
| *Elymus spicatus* | 3 | 5.689 |
| *Elymus spicatus* | 3 | 6.483 |
| *Elymus spicatus* | 3 | 8.411 |
| *Elymus spicatus* | 3 | 8.96 |
| *Elymus spicatus* | 3 | 11.327 |
| *Elymus spicatus* | 3 | 11.573 |
| *Elymus trachycaulus* | 3 | 5.756 |
| *Elymus trachycaulus* | 3 | 7.827 |
| *Elymus trachycaulus* | 3 | 8.036 |
| *Elymus trachycaulus* | 3 | 8.149 |
| *Elymus trachycaulus* | 3 | 8.905 |
| *Elymus trachycaulus* | 3 | 10.284 |
| *Elymus trachycaulus* | 3 | 11.222 |
| *Leymus cinereus* | 3 | 5.063 |
| *Leymus cinereus* | 3 | 6.071 |
| *Leymus cinereus* | 3 | 6.992 |
| *Leymus cinereus* | 3 | 7.003 |
| *Leymus cinereus* | 3 | 7.298 |
| *Leymus cinereus* | 3 | 7.816 |
| *Leymus cinereus* | 3 | 8.609 |
| *Leymus cinereus* | 3 | 17.081 |
| *Leymus condensatus* | 3 | 8.574 |
| *Leymus condensatus* | 3 | 8.685 |
| *Leymus condensatus* | 3 | 8.821 |
| *Leymus condensatus* | 3 | 9.298 |
| *Leymus condensatus* | 3 | 9.317 |
| *Leymus condensatus* | 3 | 11.011 |
| *Leymus condensatus* | 3 | 14.507 |
| *Leymus condensatus* | 3 | 14.919 |
| *Leymus triticoides* | 3 | 6.621 |
| *Leymus triticoides* | 3 | 7.18 |
| *Leymus triticoides* | 3 | 7.184 |
| *Leymus triticoides* | 3 | 7.341 |
| *Leymus triticoides* | 3 | 7.505 |
| *Leymus triticoides* | 3 | 10.96 |
| *Leymus triticoides* | 3 | 11.215 |

**Appendix 2**: Individual floret area measurements for 21 *Elymus* species and replicates.
